# Multiscale label-free volumetric holographic histopathology of thick-tissue slides with subcellular resolution

**DOI:** 10.1101/2020.07.15.205633

**Authors:** Herve Hugonnet, Yeon Wook Kim, Moosung Lee, Seungwoo Shin, Ralph H. Hruban, Seung-Mo Hong, YongKeun Park

## Abstract

Histopathology relies upon the staining and sectioning of biological tissues, which can be laborious and may cause artefacts and distort tissues. Here, we demonstrate label-free volumetric imaging of thick-tissue slides, exploiting refractive index distributions as intrinsic imaging contrast. The present method systematically exploits label-free quantitative phase imaging techniques, volumetric reconstruction of intrinsic refractive index distributions in tissues, and numerical algorithms for the seamless stitching of multiple 3D tomograms and for reducing scattering-induced image distortion. We demonstrate demonstrated label-free volumetric imaging of thick tissues with the field of view of 2 mm × 1.75 mm × 0.2 mm with a spatial resolution of 170 nm × 170 nm × 1200 nm. The number of optical modes, calculated as the reconstructed volume divided by the size of the point spread function, was approximately 20 Giga voxels. We have also demonstrated that different tumour types, and a variety of precursor lesions and pathologies can be visualized with the present method.

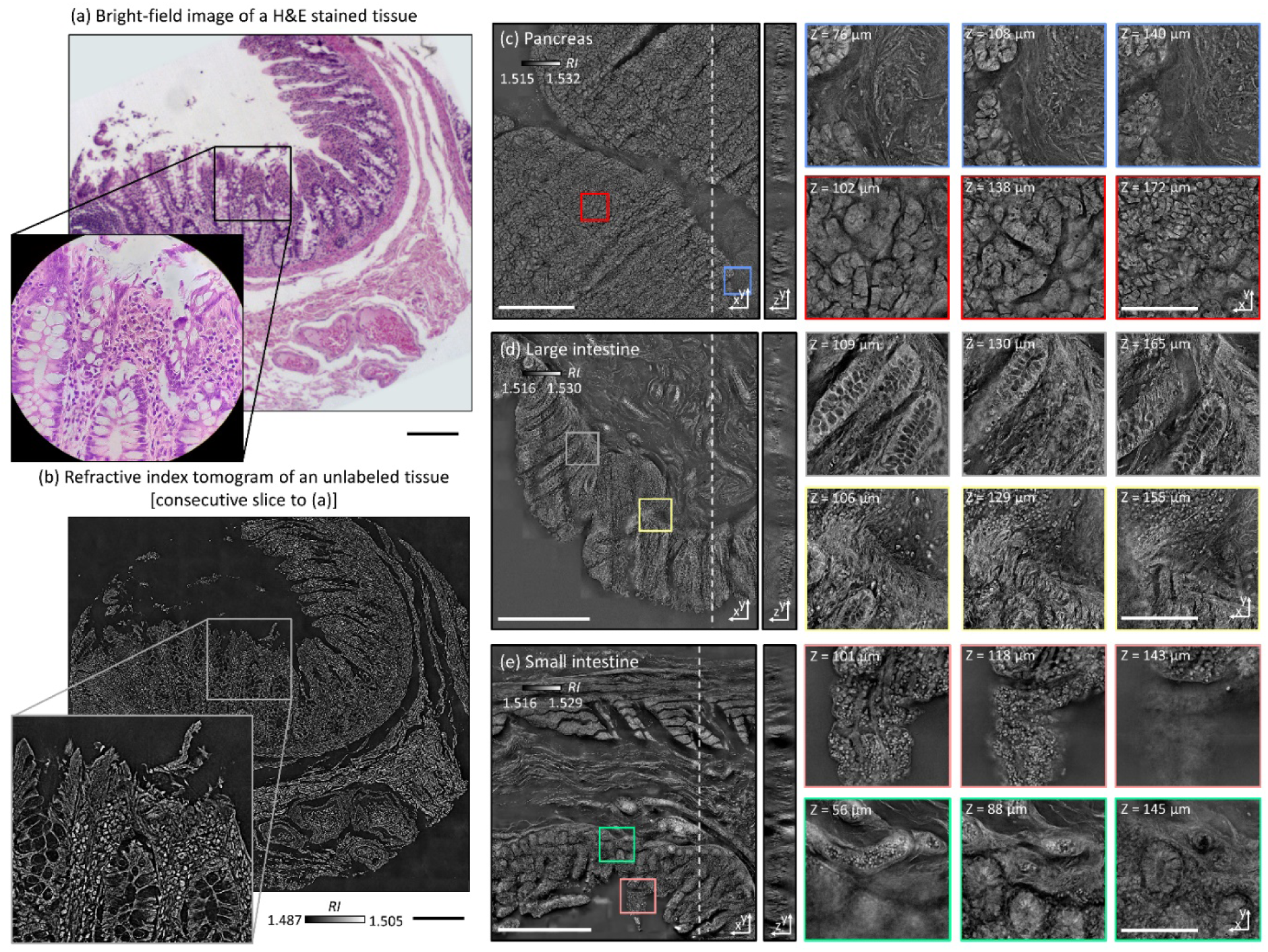

## Introduction

Microscopic assessment of biopsied and resected tissues is central to understanding the underlying pathophysiology and clinical states of many patients. Because most microscopic specimens are translucent to visible light, conventional histological methods achieve image contrast using exogenous labels, such as haematoxylin and eosin (H&E) or immunohistochemical labeling^1, 2, 3^. Although these procedures allow pathologists to easily identify cellular processes, they have several limitations, including heavy use of chemicals during sample preparation, which can induce artefacts and distort samples^4^; long sample preparation time, which can delay pathological diagnosis; and labour-intensive histochemical processes^5^. Importantly, the staining and visualization of thicker (>5 μm) tissues is limited by strong light absorption and scattering.

A variety of diverse label-free microscopy methods have been employed to circumvent these challenges. Nonlinear microscopy methods are widely used^6^ because chemical specificities are retained without exogenous labels via characteristic vibrational modes^7, 8^ or nonlinear susceptibilities^9, 10, 11, 12^ of biomolecules. Nevertheless, the long dwell time to collect weak nonlinear signals impedes high-speed image scanning and rapid identification of pathological regions. Quantitative phase imaging (QPI) is an alternative candidate for next-generation label-free histological methods because it accelerates imaging speed by exploiting holographic imaging and using elastically scattering light information as an intrinsic contrast agent for biological specimens^13^. Nevertheless, conventional QPI techniques provide only 2D information of thin tissue samples^14, 15^.

Optical diffraction tomography (ODT) is a 3D QPI technique that achieves label-free volumetric histopathology. From transmitted-light holographic measurements at various angles, similar to X-ray computed tomography, ODT reconstructs the 3D refractive index (RI) distribution of biological cells^16, 17^. Because RI values depend on the amount of intracellular biomolecules, including proteins and lipids, ODT allows label-free quantitative 3D morphological mapping of biological specimens and has been widely utilised to advance our understanding of the physiology of various live cells^18, 19, 20^. However, several technical issues, such as a small holographic field retrieval field of view (FoV) and image degradation due to multiple light scattering, have hindered 3D RI-based histopathological analyses. When using this algorithm, tissue slices with thickness of up 100 μm could be reconstructed with high contrast.

Here, we demonstrate multiscale label-free volumetric histopathology of thick tissue slides. We addressed the issues of limited FoV and image degradation by constructing a long-working-distance ODT optical setup and developing a robust multiscale ODT reconstruction and stitching algorithm, which considers optical aberration due to thick tissues. The experimental setup enabled mesoscopic imaging of human pancreatic and small and large intestine tissues over a milimetre-scale FoV with sub-micrometre resolution.

## Results

### Optical setup

To image thick tissues, a custom ODT setup was built (Fig. 1, Methods). The system was based on Mach-Zehnder interferometer, which was equipped with a long-working-distance objective lens for imaging thick tissues, an automated sample stage for raster-scanning, and a digital micromirror device (DMD) for illumination beam control. A blue continuous-wave laser (λ = 457 nm) was selected to avoid the H&E staining absorption peak. A plane wave illuminated the samples with a specific illumination angle, which was systematically controlled by projecting time-multiplexed hologram patterns on the DMD^21^. The scattered field from the sample was collected by the objective lens and projected to the camera plane where the scattered field from the sample interfered with a reference beam and generated a spatially modulated interferogram, from which both amplitude and phase images, as a function of illumination angles, were retrieved using Fourier analysis^22^ (Fig. 1c). These multiple 2D field images were used to reconstruct a 3D RI tomogram of the sample using the Fourier diffraction theorem^23^. The theoretical resolution of our imaging system was 0.17 and 1.4 μm in the lateral and axial directions, respectively^24^.

**Figure 1.**
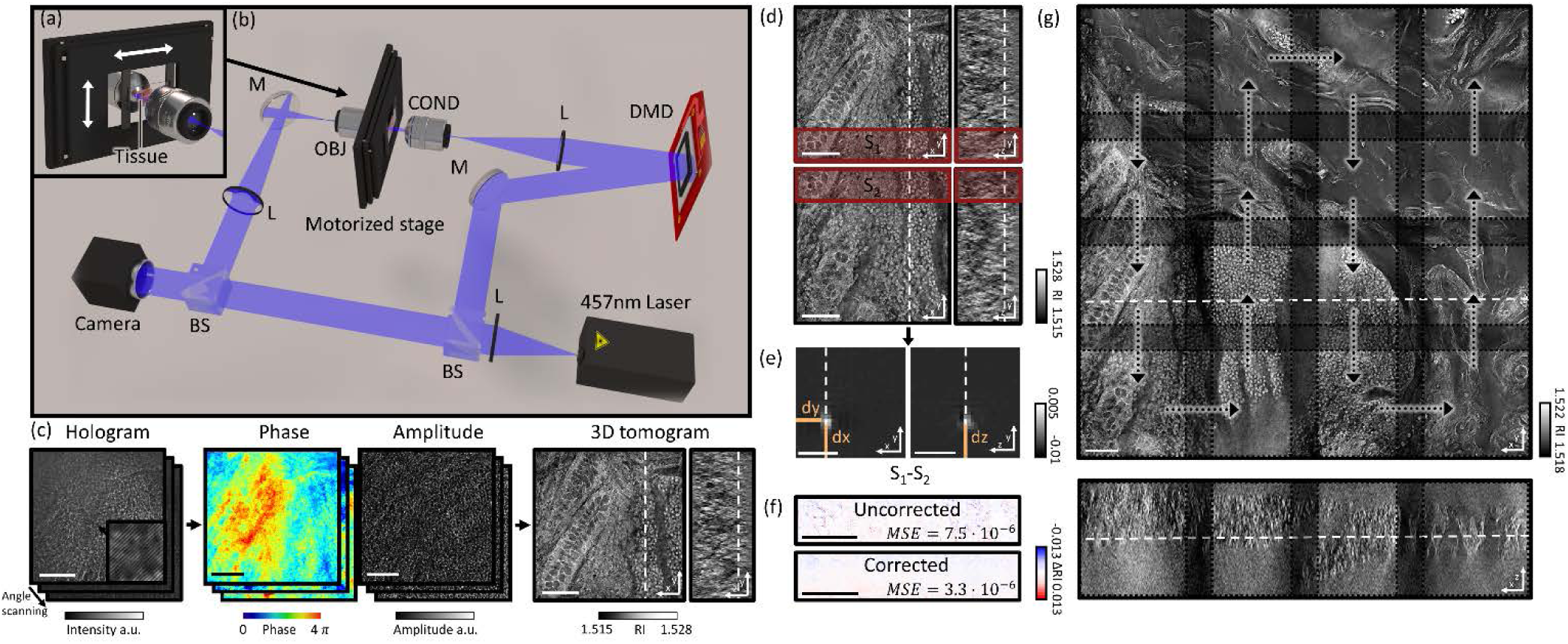
Design and mechanism of a custom optical diffraction tomography apparatus. (a) Close-up view of motorised stage; (b) Schematic of the optical setup. BS: beam-splitter; L: lens; M: mirror; DMD: digital micromirror device; COND: condenser lens; OBJ: objective lens (c) Optical diffraction tomography steps: First, a hologram is retrieved by the camera; the amplitude and phase of the field transmitted through the sample are retrieved from this hologram. Finally, the refractive index map of the sample is obtained. (d–g) Stitching algorithm steps: (d) Overlapping regions of adjacent tiles are retrieved; (e) Phase correlation algorithm output allows retrieval of the sub-pixel shift between the two overlapping regions; (f) Mean square error between the overlapping regions after the subpixel shift correction; (g) Small fields of view are raster scanned to form a big field of view; they are then stitched together using the relative position found in (e)

There are two main technical challenges in the reconstruction of wide and thick biological tissues: (1) alignment of multiple overlapping image tiles into a single stitched image, and (2) improvement of image contrast against sample-induced aberrations. We addressed these issues computationally using correlation-based subpixel adjustments and digital field refocusing during RI reconstruction, respectively.

### Stitching segmented tomograms

First, accurate image alignment between overlapping image tiles was achieved by correlation-based subpixel adjustment (Figs. 1d–g). During data acquisition, overlapping regions between adjacent 3D tiles was set to 20%, which guaranteed border artefact suppression during RI reconstruction (Fig. 1g). The precise relative 3D position between overlapping regions was then determined with sub-pixel accuracy using a phase-correlation algorithm^25^ (Fig. 1e). This position-correcting process enabled artefact reduction due to image misalignment, as evidenced by a mean squared errors (MSEs) analysis (Fig. 1f). Next, the global positions of multiple image tiles were further adjusted using least square minimisation^26^. Minimisation was weighted using the Pearson correlation coefficient of overlapping sections. This allowed quantifying similarity between overlapping regions and thus gave less importance to regions without tissue where finding the relative position was more difficult. Weighting and the use of robust least square minimisation avoided error propagation. Finally, images were placed at the obtained position and seamlessly blended together (Fig. 1g, 4d).

An important consideration when using stitching with ODT is that the illumination and detection size of single tiles is spatially limited. This limits the maximal axial field of view. Indeed, the volume that receives the detection at full numerical aperture and that is illuminated by every illumination angle is limited to a conical region above and below the focal plane. Out of this region, the image will have both lower resolution and ghost images due to the circular symmetry of the discrete Fourier transform used in the reconstruction algorithm. It is important to overlap the tiles sufficiently, so that these erroneous regions of the tomograms are removed when stitching. In all the experiments, an overlapping distance corresponding to 20% of the camera field-of-view was used.

### Numerical focusing through thick tissues

The use of coherent light in ODT allows the focal plane to be adjusted numerical, avoiding blurring artefact due to focus mismatching, which occurs in bright-field microscopy and other incoherent imaging methods. In our reconstruction algorithm, when imaging thin samples, the sample position is first automatically detected, then an image containing only the most in-focus part of the tissue is generated. The optimal focus plane was found by using the Tamura of the gradient focus criterion^27^, which is a function that reaches its maximum when an image is well focused. The focal position was researched in this way every 40 μm and interpolated in regions without samples. This method was also used in thick tissue to quickly find high contrast well focused lateral cross-sections.

### Reconstruction of refractive index distributions

Initially reconstructed tomograms of thick tissues suffered from weak image contrast and distortion due to sample-induced aberrations. Considering that samples nearest the optical focal plane display sharper image contrast when reconstructed with the Rytov approximation, we exploited holographic refocusing to reconstruct 3D RI maps with higher image contrast^28, 29^ (Figs. 4a-c). We digitally refocused the obtained light field images to virtual focal planes using a diffraction kernel based on Green’s function (Fig. 4a). The refocusing interval is chosen as a compromise between the computation speed and contrast lost between the virtual focal planes. Here, we set the refocusing interval to 9 μm, reconstructed RI tomograms 9 μm thick around each virtual focal plane, and combined the tomogram stack into a single tomogram. Qualitative comparison validated that the RI tomograms out of a focal plane could recover image contrast using refocusing-based RI reconstruction (Fig. 4c). The RI distribution root-mean-squares also increased at the out-of-focus planes, quantitatively confirming that digital refocusing improved RI contrast of reconstructed thick tissue tomograms (Fig. 4c).

These algorithms were implemented on a graphical processing unit (GPU) to facilitate reconstruction speed. In particular, since phase unwrapping has to be performed at every refocusing steps, phase unwrapping is a particularly computationally intensive part of the process. To improve performance we implemented a custom version of Goldstein’s phase unwrapping algorithm^30^ where the residue pairing was realise on the CPU while residue detection, residue linking, rasterization and phase unwrapping where performed on the GPU.

### Performance in thick tissues

To validate the capacity of the present method for tissue slides volumetric histopathology, we first measured human small intestine tissue samples. For comparison purposes, we prepared two consecutive tissue slides, one using conventional H&E staining and the other unlabelled. The XY cross-sectional images of the unlabelled slide in the optical focal plane were consistent with the conventional in-focus bright-field images of the H&E stained slide, demonstrating the high accuracy of this method (Figs. 2a–b). Overall tissue anatomy and subcellular features were clearly seen in both the conventional H&E image and RI tomogram. To demonstrate applicability, various histopathology tissue slides were imaged (Figs. 2c–e). Unlabelled tissue samples from human organs (pancreas, small intestines, and large intestines) were cut at a thickness of 100 μm and analysed using the present method. The resulting FoVs were 2 mm × 1.75 mm x 0.2 mm. The capability of volumetric histopathology can be seen in the high-resolution images of subparts at various axial foci (insets, Figs. 2c-e). Not only can subcellular features be accessed without labelling, but the three-dimensional architecture of tissue structures can also be investigated.

**Figure 2.**
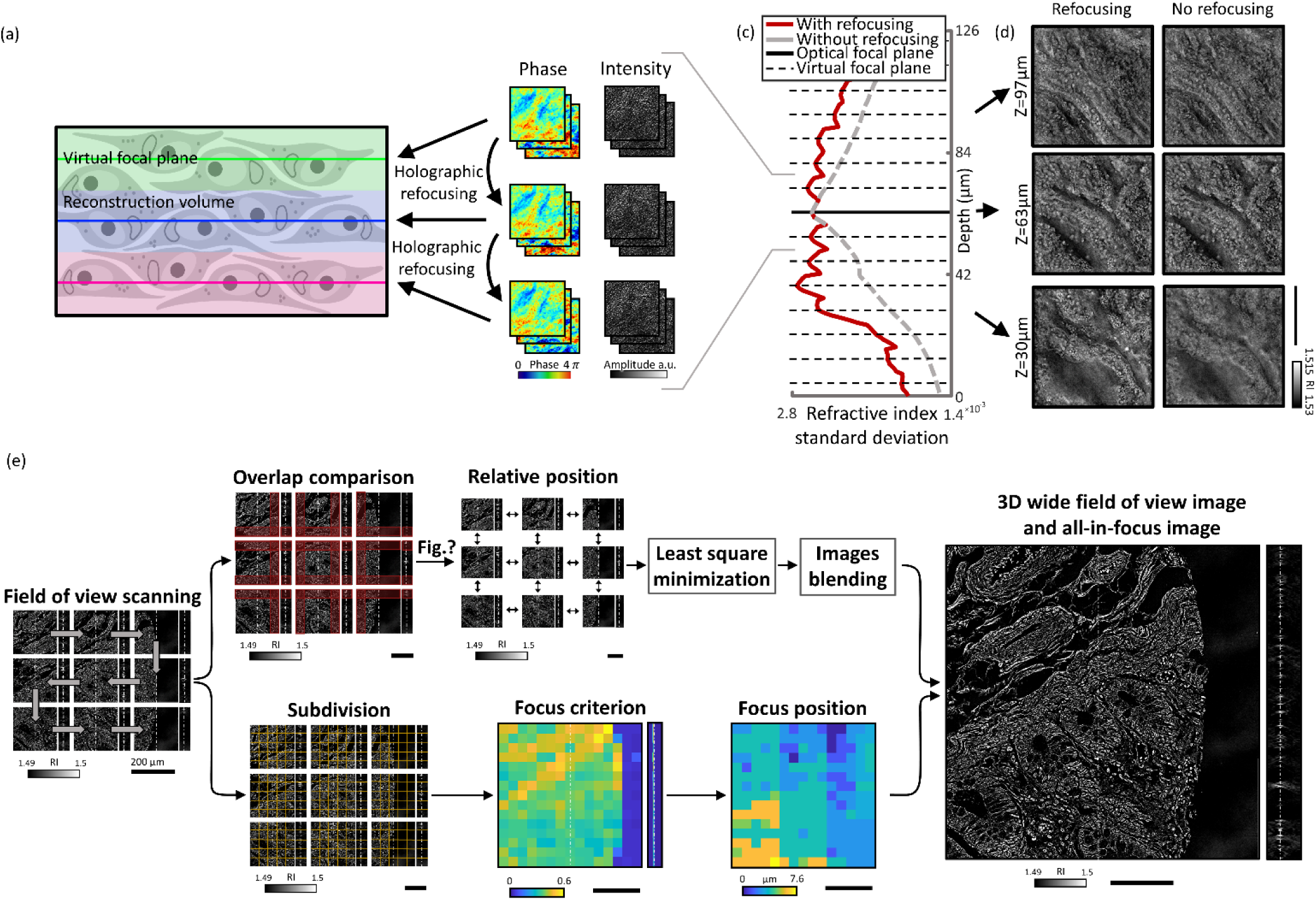
(a) Schematic of virtual focal planes and holographic refocusing; (b) RI standard deviation as a function of physical depth; (c) Cross-sectional images of the reconstructed tomogram (left) with holographic refocusing and (right) without holographic refocusing (d) Schematic of the stitching and focus finding steps. Scale bars are 100 μm long.

### Imaging pathologic tissue slides

To validate further the present method for clinical applications, volumetric images of unlabelled thick pancreas tissues samples obtained from patients with pancreas neoplasms are shown in Fig. 3. The RI tomograms of tissues from patient #1 (Fig. 3a) shows neoplastic epithelial cells with phenotypic neuroendocrine differentiation, characteristics of a pancreatic neuroendocrine tumour (PanNET). The back-to back neoplastic cells with minimal stroma are easily appreciated, as is the uniformity in the shape of the round nuclei. All of these allow for the recognition of the diagnosis. For validation purposes, adjacent tissues were prepared and images using the conventional H&E staining method, which exhibit good agreement with the present method. The RI tomograms of the tissues from patient #2 show low grade pancreatic intraepithelial neoplasia (PanIN) as well as normal pancreatic ducts, which show different cytoplasmic features between normal pancreatic duct and low grade PanIN. In the liver tissues from patient #3, normal bile duct, intraductal papillary neoplasm of bile duct (IPNB), and cirrhotic nodule are imaged with the present method. Thus, different tumour types, and a variety of precursor lesions and pathologies can be visualized.

**Figure 3.**
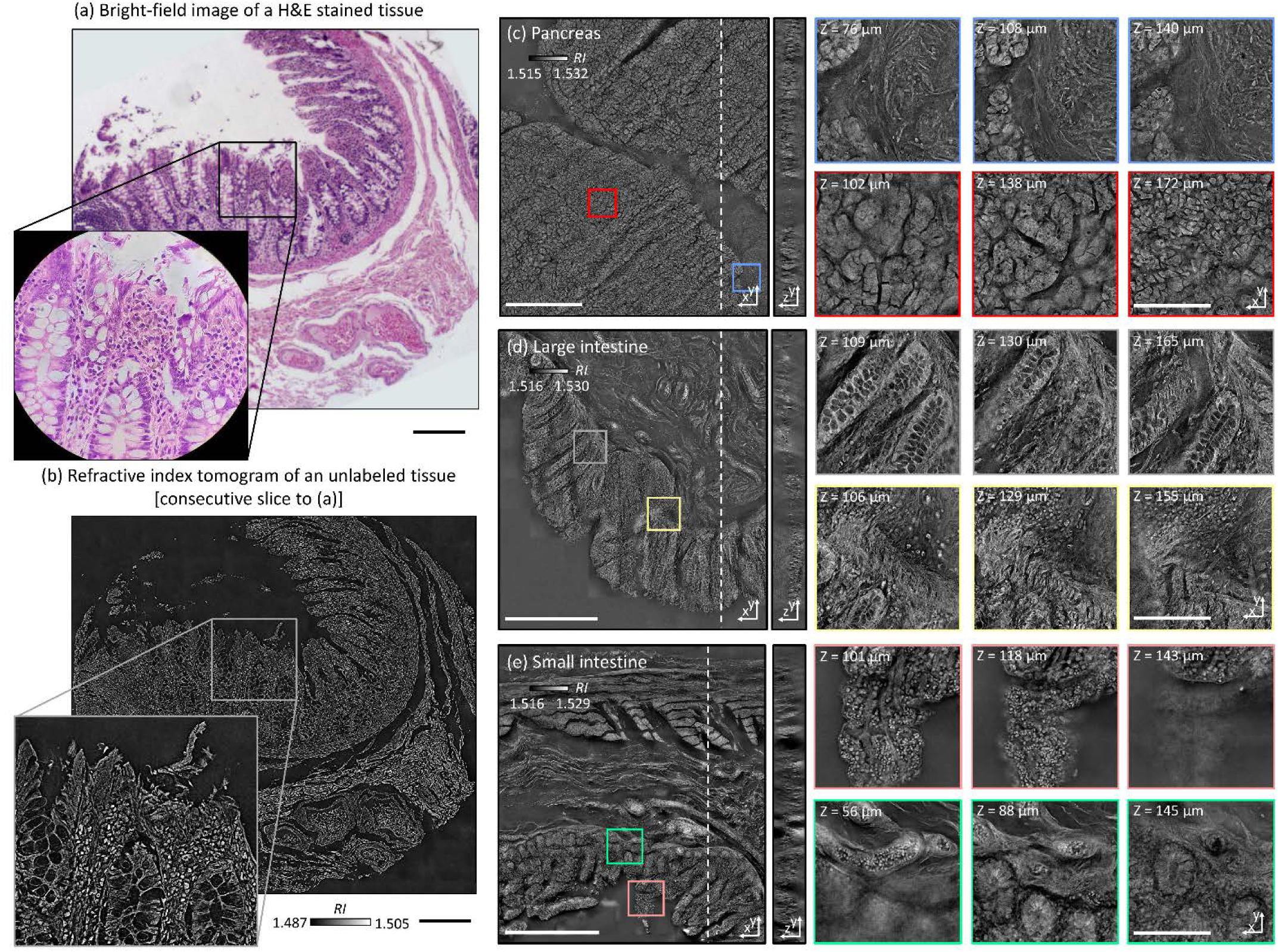
Comparison of a normal colon tissue slice imaged with bright field and stained with H&E staining (a) and an unstained neighbor tissue slice imaged with ODT (b). In (a), the zoomed in image was taken with a 60× 0.8 NA objective lens, while in (b), the zoomed in image is a cropped version of the stitched image. (c-e) 3D wide field images of 100-μm-thick tissue slices of (c) pancreas, (d) colon, and (e) small intestine. Zoomed in regions of interest are shown at different depths. Scale bars are 100 μm long in (a-b) and zoom in from (c-e). Scale bars are 500 μm long in in wide field of view images (c-e).

**Figure 4.**
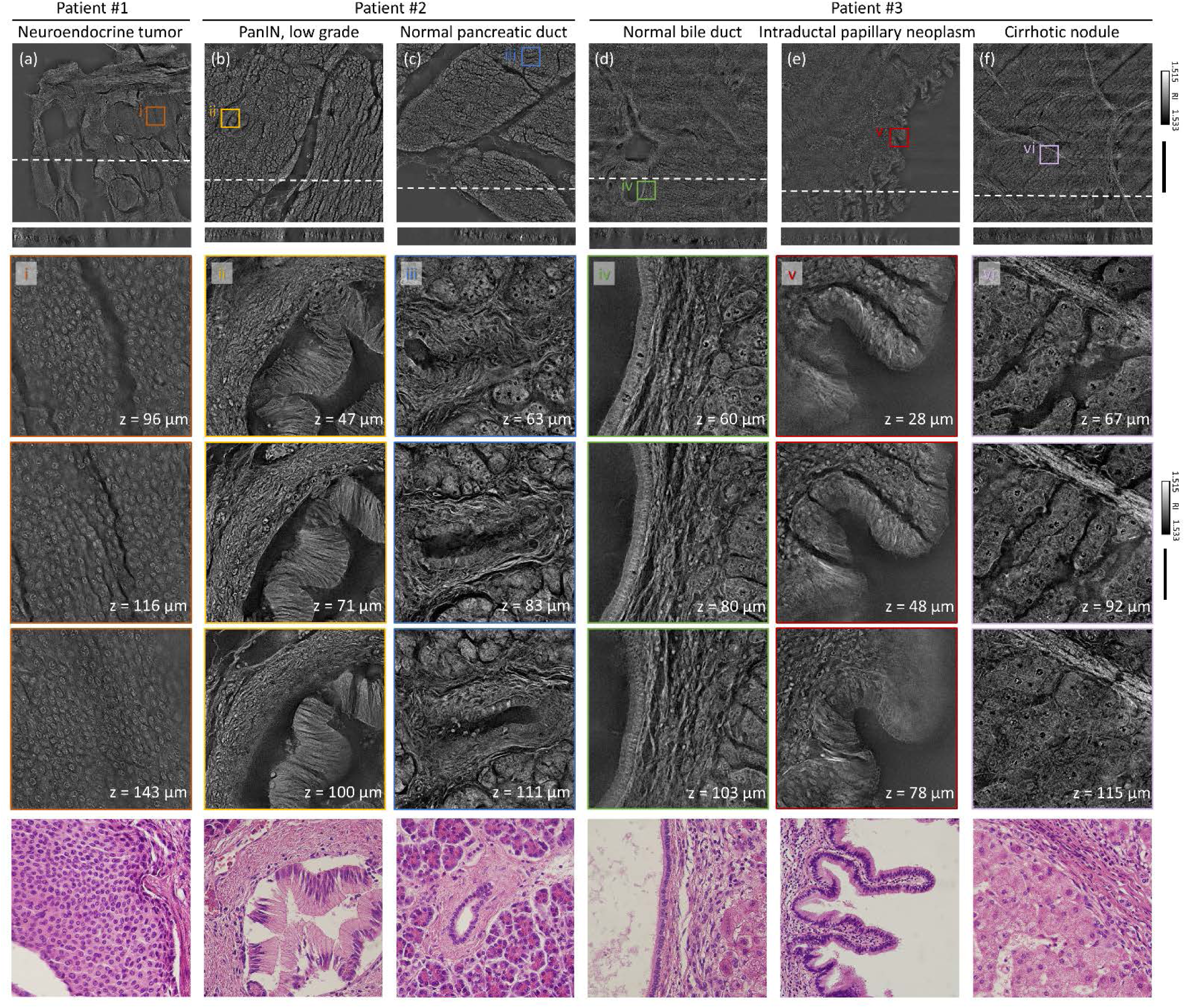
Volumetric histopathoogy of unlabeled 150-μm-thick pancrea tissue samples from three individuals: (a) patient with a pancreatic neuroendocrine tumor; (b) patient with pancreatic intraepithelial neoplasia (PanIN); and (c) patient with intraductal papillary neoplasm of bile duct in the liver. The areas indicated with the boxes i – vi are shown at three different axial positions with manification (from the second to the forth rows). For comparision purpose, adjacent tissues were prepared in thin tissue slides with conventional H&E staining method. (the fifth row, 400x magnification). Scale bars are 500 μm long in wide field of view images and 50 μm long in zoomed regions of interest.

## Discussion

In summary, we present multiscale volumetric imaging of unlabelled tissue samples. We imaged millimetre-scale, unstained, 100-μm-thick tissues at a subcellular three-dimensional resolution, which enabled the visualisation of individual cells and multicellular tissue architectures in a variety of tissues. By systematically stitching 3D RI tile data obtained with a long-working-distance ODT setup, we demonstrated a FoV of 2 mm × 1.75 mm × 0.2 mm with a spatial resolution of 170 nm × 170 nm × 1200 nm, and throughout this extended imaging volume, image degradation due to sample-induced aberration was significantly reduced using the holographic refocusing strategy. The present method can be understood as extremely high content imaging. The number of optical modes, calculated as the reconstructed volume divided by the size of the point spread function, was approximately 20 Giga voxels.

Considering the label-free volumetric imaging capability exhibited across a variety of tissue samples, this method could potentially be used for rapid cancer diagnosis during intraoperative pathology consultations or small biopsy samples. In addition, the present digital pathology approach has considerable advantages over conventional imaging methods, particularly in terms of measuring seamless multiscale volumetric images; conventional methods only generate one or a few focused plane images. In the current system, the time needed to measure 11 × 11 sections (1.75 mm × 1.75 mm) and produce volumetric reconstructions is 40 minutes and 135 minutes, respectively. However, this can be further expedited by implementing a high-speed translational stage, a fast image sensor, and more powerful computing power.

To make the method more applicable, more research is needed on the histological interpretation of RI information. Because RI tomograms do not generate the same data as H&E staining, they would be utilised as complementary information. We envision that this generic approach could have far-reaching applications in histopathology and cytometry, possibly in conjunction with newly emerging machine learning methods for segmentation and cell-type classification.

## Methods

### Materials and Methods

#### Long working-distance optical diffraction tomography setup

A schematic image of the optical setup is presented in Fig. 1(a). A blue continuous-wave laser (central wavelength = 457 nm, Cobolt Twist, Cobolt) was used as a coherent illumination source. For ODT, a Mach-Zehnder interferometer was installed with a digital micromirror device (DMD; DLPLCR6500EVM, Texas Instrument) in the sample arm for high-speed illumination scanning^21, 31^. The angled plane wave diffracted from the DMD was demagnified by a *4*-*f* lens array, including a long-working-distance condenser objective (LUCPLFLN40X, numerical aperture (NA) = 0.6, Olympus). The scattered field from the tissue was collected using an objective lens (UPlanSAPO20X, NA = 0.75, Olympus), magnified by a factor 55, interfered with the reference beam, and imaged with a CMOS camera (LT425M-WOCG, Lumenera Inc.). For holographic field retrieval, the resulting interferograms were converted to amplitude and phase images using a field retrieval algorithm based on Fourier analysis^22^ (Fig. 1c). The retrieved field images were used to reconstruct a 3D RI tomogram of the tissue using the Fourier diffraction theorem^23^.

#### Holographic refocusing of tissue slides

The use of coherent light in ODT allows the focal plane to be adjusted numerical, avoiding blurring artefact due to focus mismatching, which occurs in bright-field microscopy and other incoherent imaging methods. In our reconstruction algorithm, when imaging thin samples, the sample position is first automatically detected, then an image containing only the most in-focus part of the tissue is generated. The optimal focus plane was found by using the Tamura of the gradient focus criterion^27^, which is a function that reaches its maximum when an image is well focused. The focal position was researched in this way every 40 μm and interpolated in regions without samples.

#### Evaluation of optical diffraction tomography in thick samples

Refractive index reconstruction using the Fourier diffraction theorem^23^ is based on a weak scattering approximation and as such does not hold well as samples become thicker. Recently different methods mitigating multiple scattering effects on image reconstruction have been developed^32, 33, 34^. However, these are computationally expansive, and are still not suitable for a wide field of view imaging. However, it has also been shown that refocusing the field at every focal plane before reconstructing the refractive index can increase contrast in thick samples^28, 29^. This method is less efficient than the previous ones, but it is computationally less expansive and has been used in our algorithm.

Furthermore, we only refocused the field at a few focal planes and then reconstructed the refractive index around those virtual focal planes. This decreases the computational cost while conserving the quality improvement from the original refocusing method. To estimate the efficiency of our algorithm, the same data were reconstructed with and without digital refocusing. Then the imaging quality was assessed by computing the refractive index contrast using its standard deviation. Indeed, sample induced aberrations cause the fields to add up destructively, reducing both contrast and resolution. From Figs. 1(i)-(j), one can see that when digital refocusing was not used, the contrast rapidly decreases from the optical focal plane. However, when refocusing is used, contrast again increases at every virtual focal plane. The refocusing interval is chosen as a compromise between the computation speed and contrast lost between the virtual focal planes. In this experiment, a refocussing interval of 9 μm was used. Even when using digital refocusing, due to multiple light scattering, the quality of the retrieved tomograms degraded as the tissue thickness was increased. Due to this, the tissue thickness is limited to approximately 100 μm when using the current setup and sample preparation.

A last limiting factor is that the illumination and detection size of single tiles is spatially limited. This limits the maximal axial field of view. Indeed, the volume that receives the detection at full numerical aperture and that is illuminated by every illumination angle is limited to a conical region above and below the focal plane. Out of this region, the image will have both lower resolution and ghost images due to the circular symmetry of the discrete Fourier transform used in the reconstruction algorithm. It is important to overlap the tiles sufficiently, so that these erroneous regions of the tomograms are removed when stitching. In all the experiments, an overlapping distance corresponding to 20% of the camera field-of-view was used.

#### Sample preparation

It is crucial when imaging thick samples to use RI matching to limit multiple light scattering. Formalin fixed paraffin embed tissue blocks from normal small intestine, colon, and pancreas, and from pancreatic neuroendocrine tumor and intraductal papillary neoplasm of the bile duct were selected. Tissue blocks were sliced with 100 μm thickness, deparaffinised using three consecutive 10 minutes xylene bath and where finally mounted between two number zero coverslips using Permount mounting medium (Permount, RI = 1.52, Fisher Chemical). Thin 3 μm thick sections were mounted on 1 mm thick slide glass with Acrymount mounting medium.

#### Ethics statement

Human studies were conducted according to the principles of the Declaration of Helsinki and were approved by the Institutional Review Board of Asan Medical Center (IRB project number: 2020-0354, Seoul, Republic of Korea) with a waiver of informed consent.

## Acknowledgements

This work was supported by KAIST, BK21+ program, Tomocube, and National Research Foundation of Korea (2017M3C1A3013923, 2015R1A3A2066550, 2018K000396).

## Author contributions

H.H., S.-M.H., and Y.P. conceived the initial idea. H.H. develop the optical system and analysis methods. H.H., XXX performed the experiments and analysed the data. M.L., S.S., YYY provided the analysis methods and analysed the data. All authors wrote and revised the manuscript. Correspondence and requests for materials should be addressed to Y.P.

## Methods

### Materials and Methods

#### Long working-distance optical diffraction tomography setup

A schematic image of the optical setup is presented in Fig. 1a-b. A blue continuous-wave laser (central wavelength = 457 nm, Cobolt Twist, Cobolt) was used as a coherent illumination source. For ODT, a Mach-Zehnder interferometer was installed with a digital micromirror device (DMD; DLPLCR6500EVM, Texas Instrument) in the sample arm for high-speed illumination scanning^21, 31^. The angled plane wave diffracted from the DMD was demagnified by a *4*-*f* lens array, including a long-working-distance condenser objective (LUCPLFLN40X, numerical aperture (NA) = 0.6, Olympus). The scattered field from the tissue was collected using an objective lens (UPlanSAPO20X, NA = 0.75, Olympus), magnified by a factor 55, interfered with the reference beam, and imaged with a CMOS camera (LT425M-WOCG, Lumenera Inc.). For holographic field retrieval, the resulting interferograms were converted to amplitude and phase images using a field retrieval algorithm based on Fourier analysis^22^ (Fig. 1c). The retrieved field images were used to reconstruct a 3D RI tomogram of the tissue using the Fourier diffraction theorem^23^.

