## Supplementary Information for "Multiscale label-free volumetric holographic histopathology of thick-tissue slides with subcellular resolution"

Supplementary Figures 1-3 | Cross-sectional image of label-free 3D wide field image of the 100 μm thick tissue slices of pancreas. This is the magnified version of the image shown in Fig. 2(c-e). The same scale and colormap used.

Supplementary Figure 4-9 | Cross-sectional image of label-free 3D wide field image of the 100 μm thick tissue slices. This is the magnified version of the image shown in Fig. 3(a-f). The same scale and colormap used.

Cross-section images in Supplementary Figs. 1-9 are available at full resolution at the figshare (https://figshare.com/account/home#/projects/78624).

cross-section imaged in d–f is available at full resolution at http://www.gigapan.com/gigapans/199300

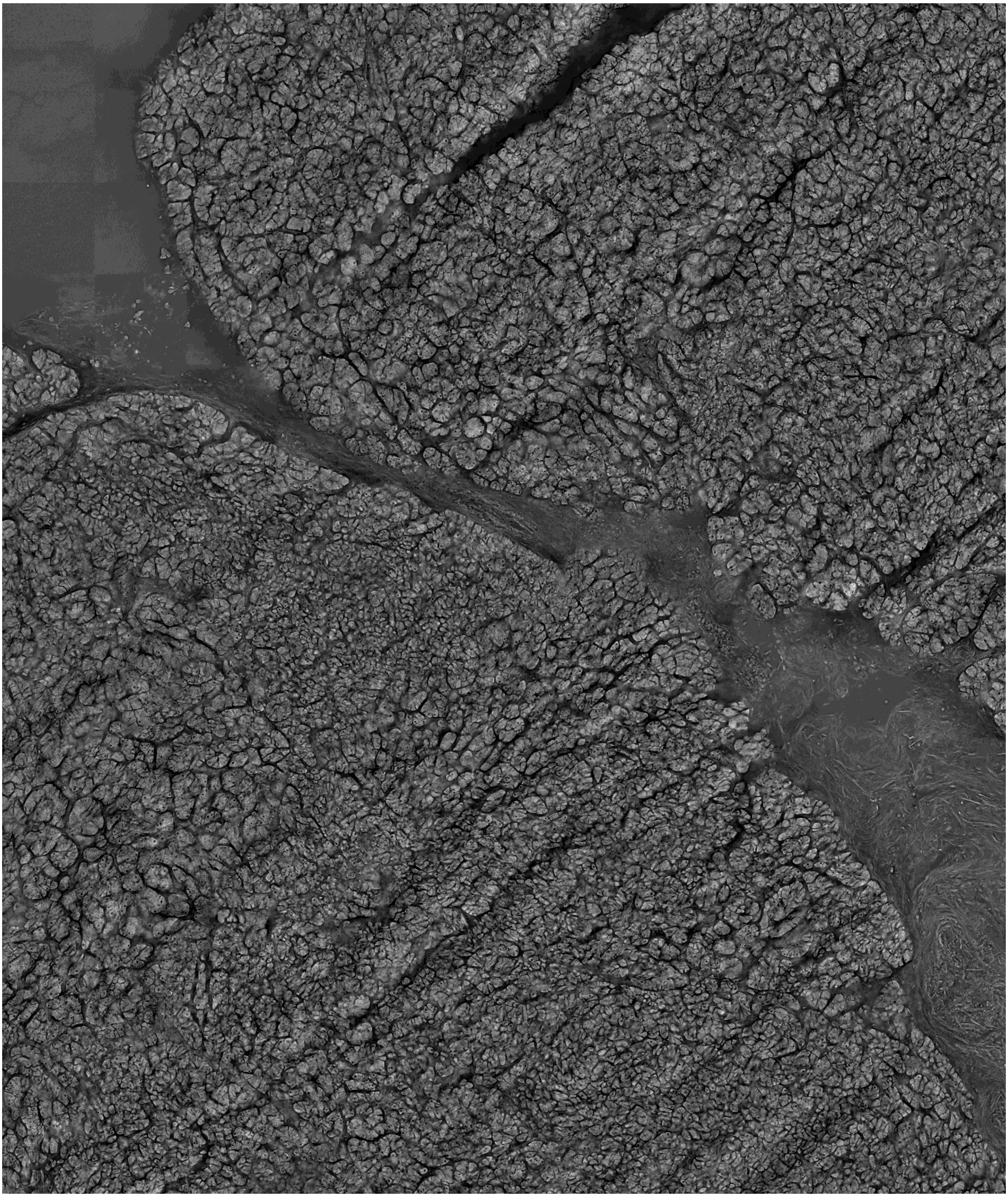

Supplementary Figure 1

Cross-sectional image of label-free 3D wide field image of the 100 μm thick tissue slices of pancreas. This is the magnified version of the image shown in Fig. 2(c). The same scale and colormap used.

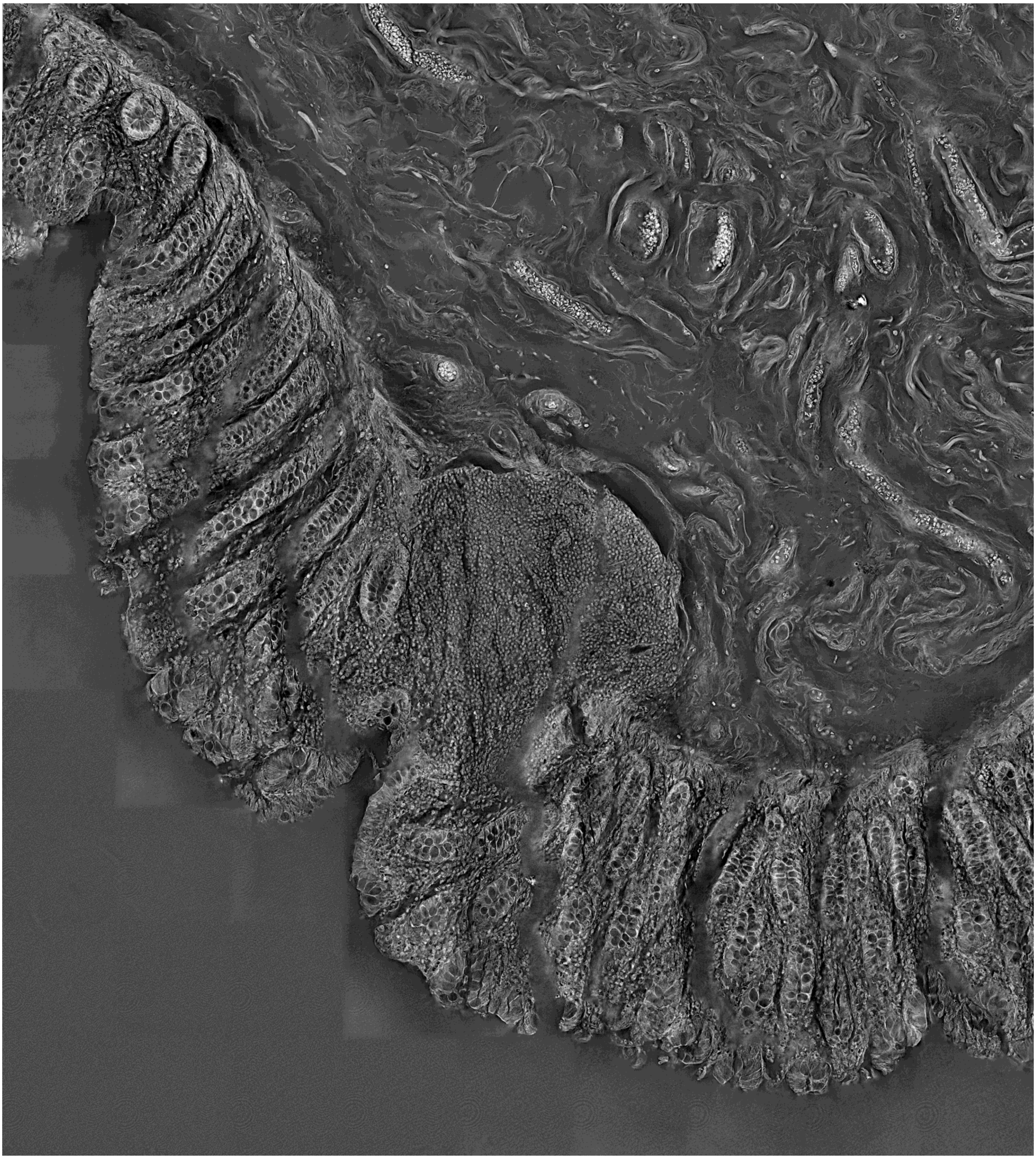

Supplementary Figure 2

Cross-sectional image of label-free 3D wide field image of the 100 μm thick tissue slices of big intestine. This is the magnified version of the image shown in Fig. 2(d). The same scale and colormap used.

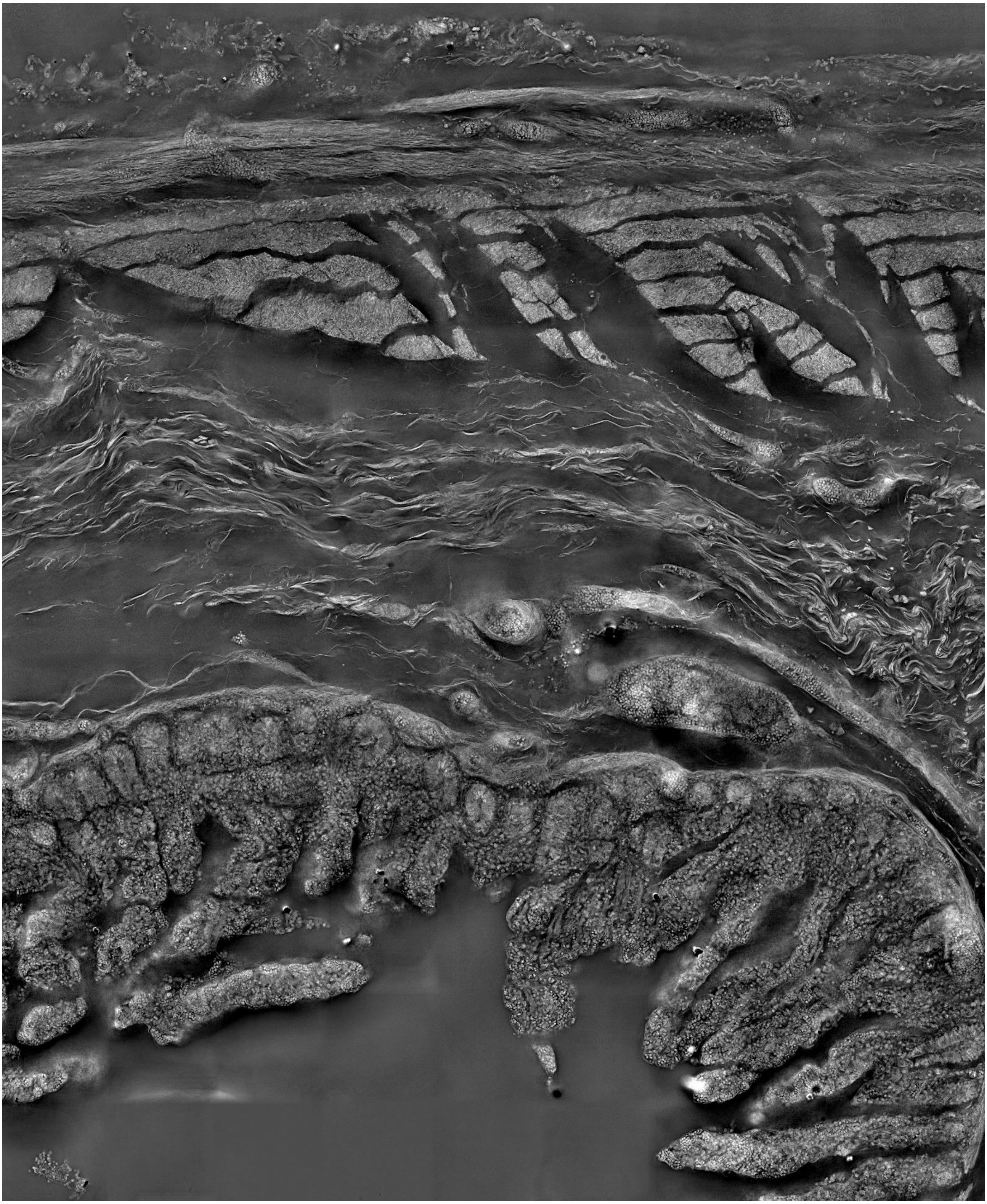

Supplementary Figure 3

Cross-sectional image of label-free 3D wide field image of the 100 μm thick tissue slices of small intestine. This is the magnified version of the image shown in Fig. 2(e). The same scale and colormap used.

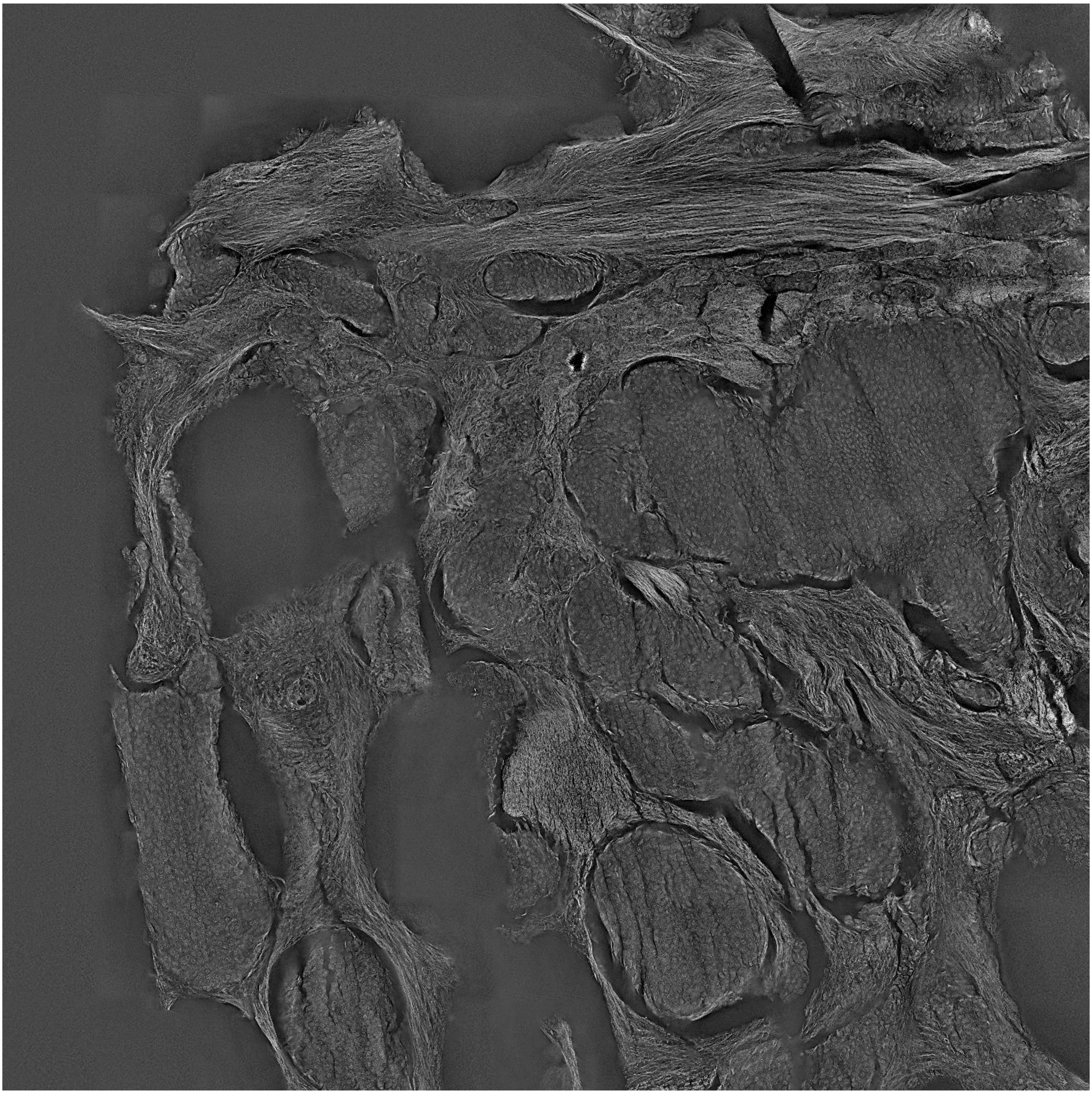

Supplementary Figure 4

Cross-sectional image of label-free 3D wide field image of the 100 μm thick tissue slices. This is the magnified version of the image shown in Fig. 3(a). The same scale and colormap used.

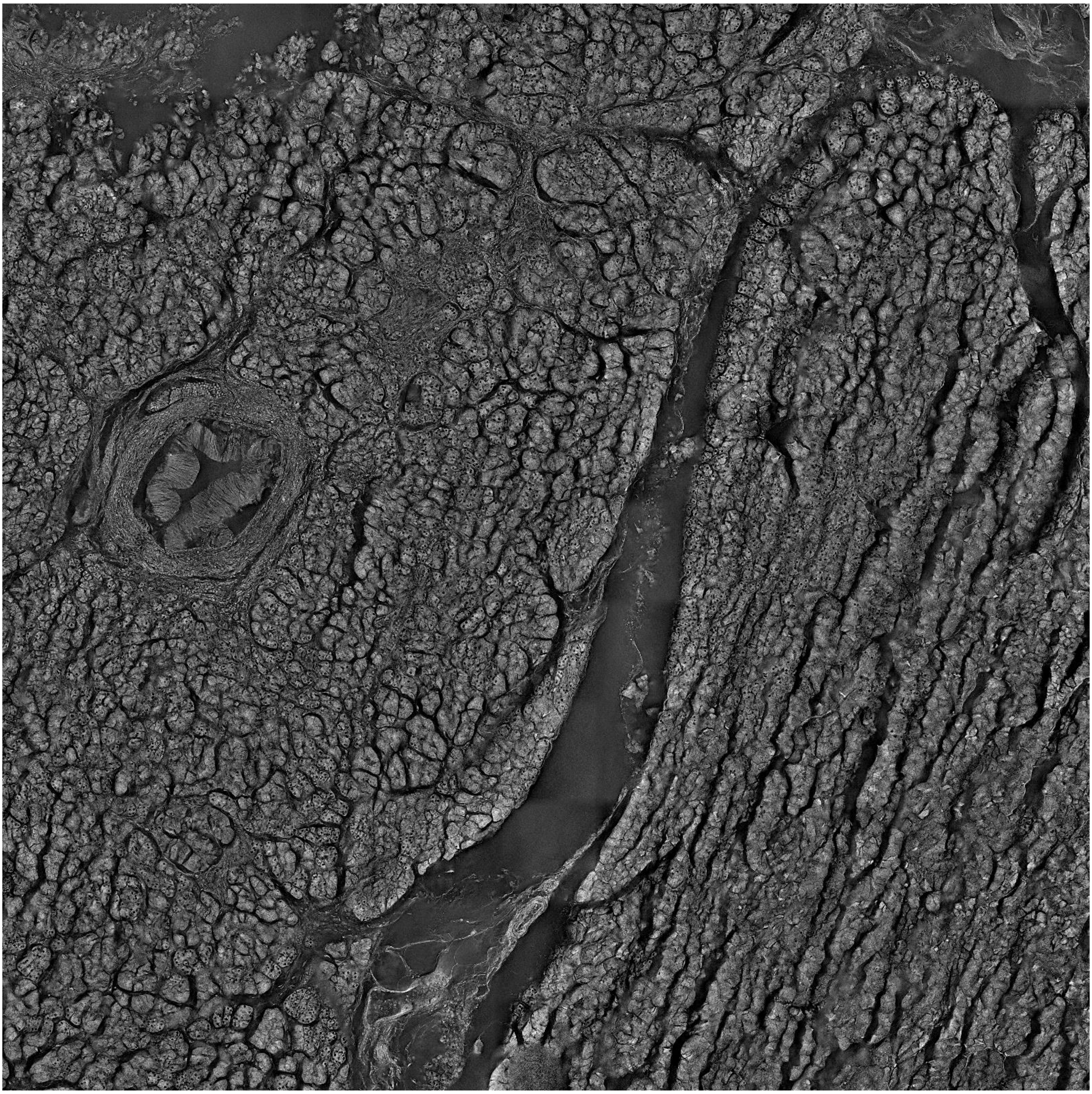

Supplementary Figure 5

Cross-sectional image of label-free 3D wide field image of the 100 μm thick tissue slices. This is the magnified version of the image shown in Fig. 3(b). The same scale and colormap used.

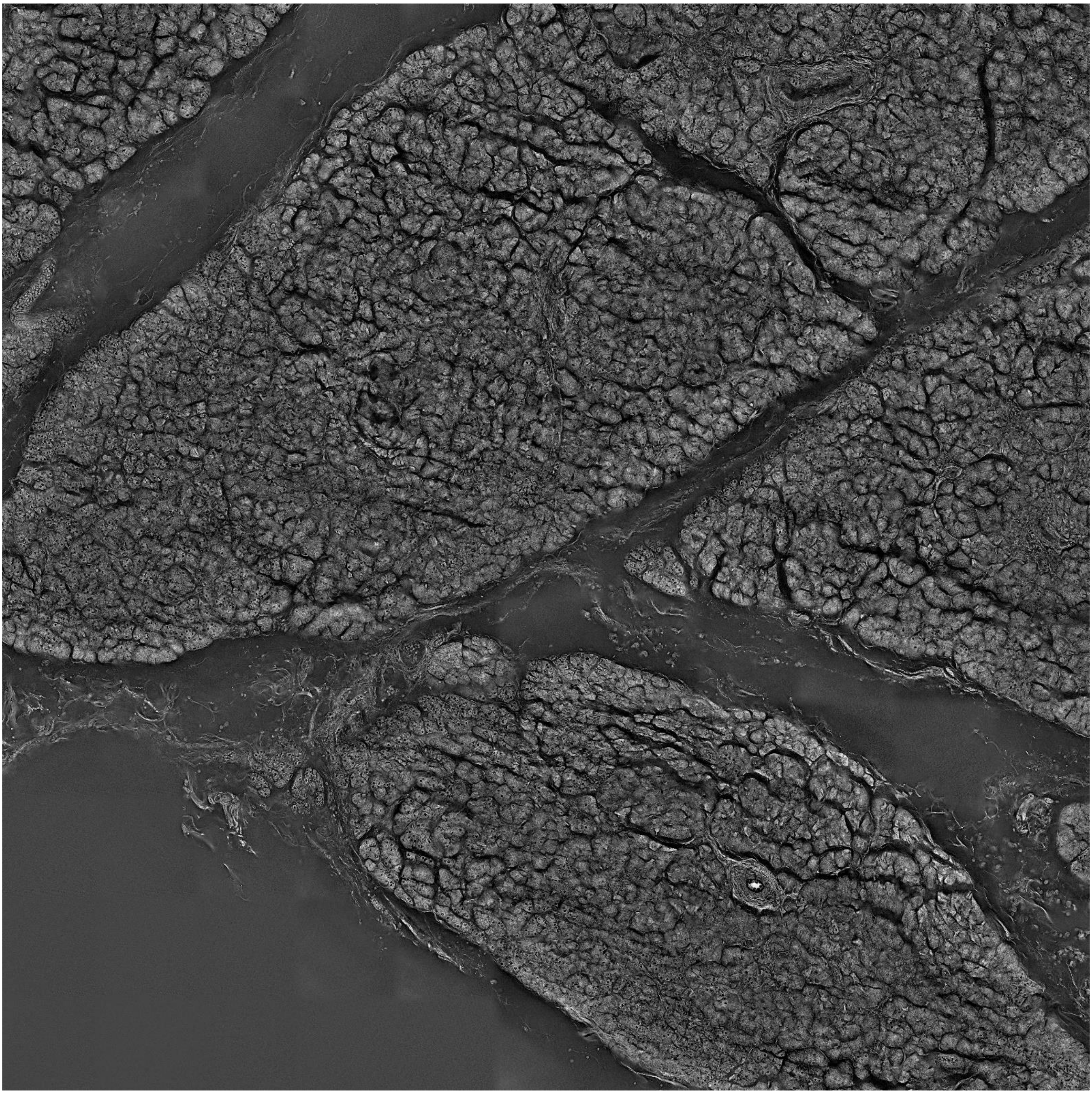

Supplementary Figure 6

Cross-sectional image of label-free 3D wide field image of the 100 μm thick tissue slices. This is the magnified version of the image shown in Fig. 3(c). The same scale and colormap used.

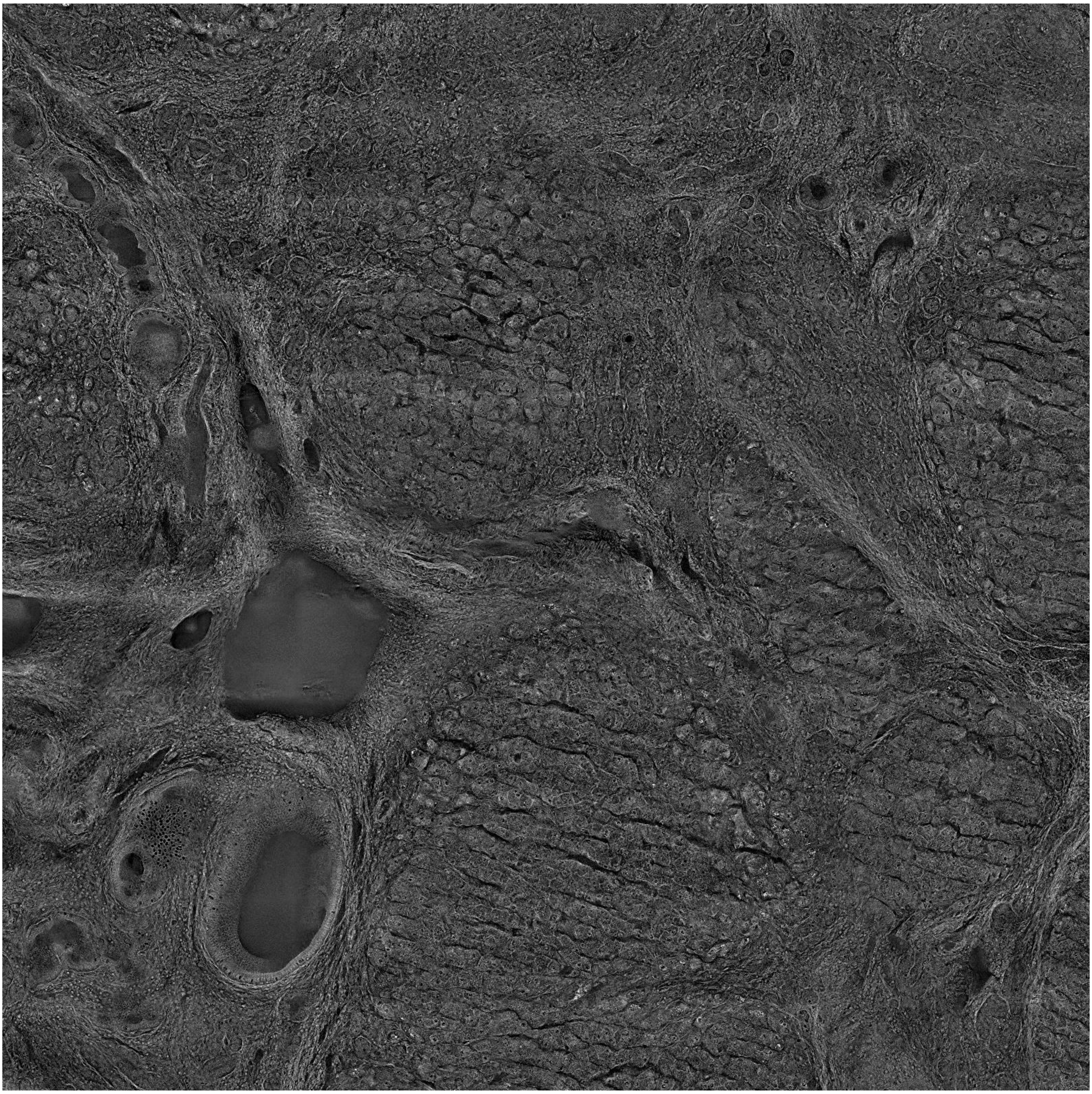

Supplementary Figure 7

Cross-sectional image of label-free 3D wide field image of the 100 μm thick tissue slices. This is the magnified version of the image shown in Fig. 3(d). The same scale and colormap used.

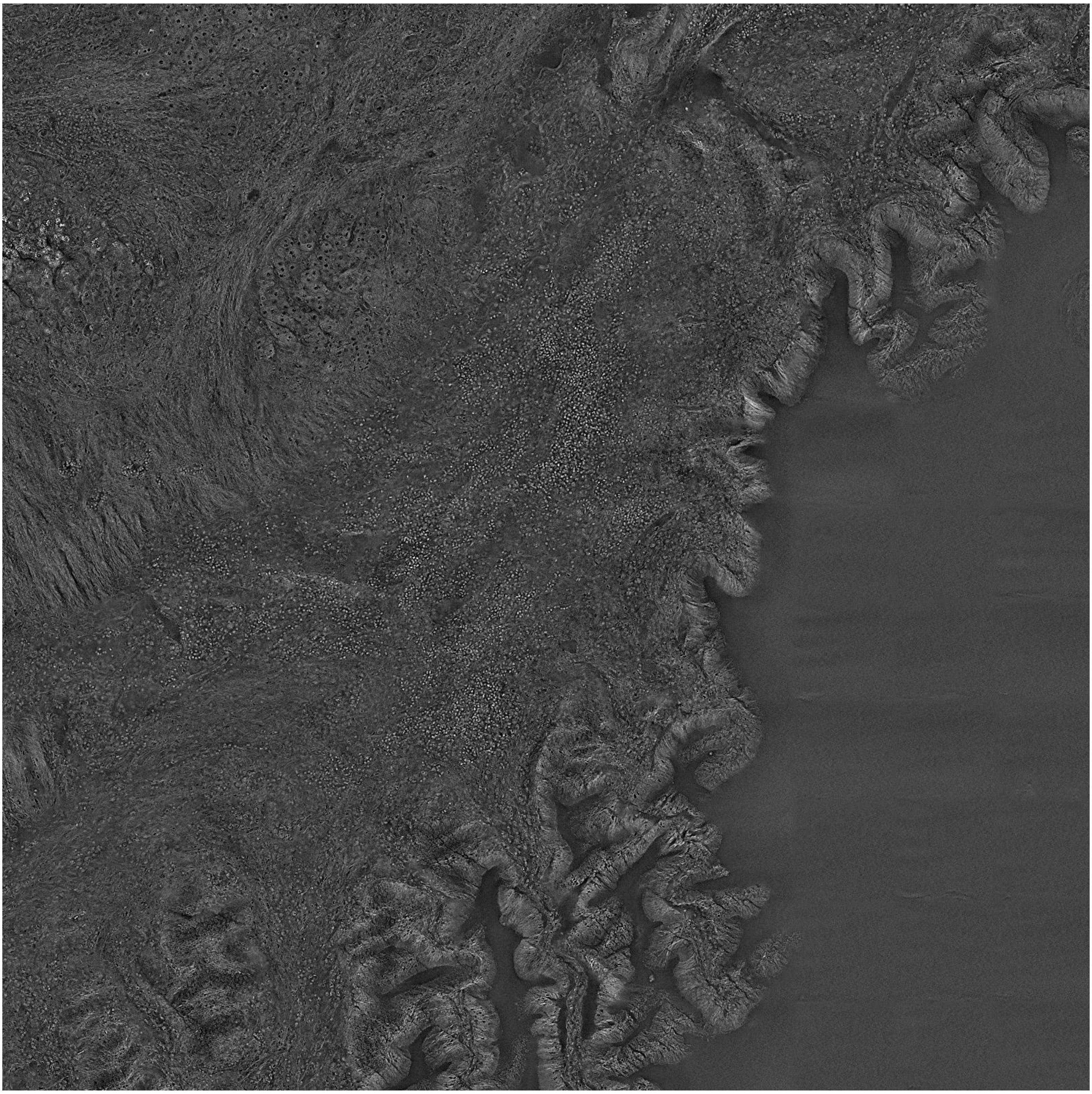

Supplementary Figure 8

Cross-sectional image of label-free 3D wide field image of the 100 μm thick tissue slices. This is the magnified version of the image shown in Fig. 3(e). The same scale and colormap used.

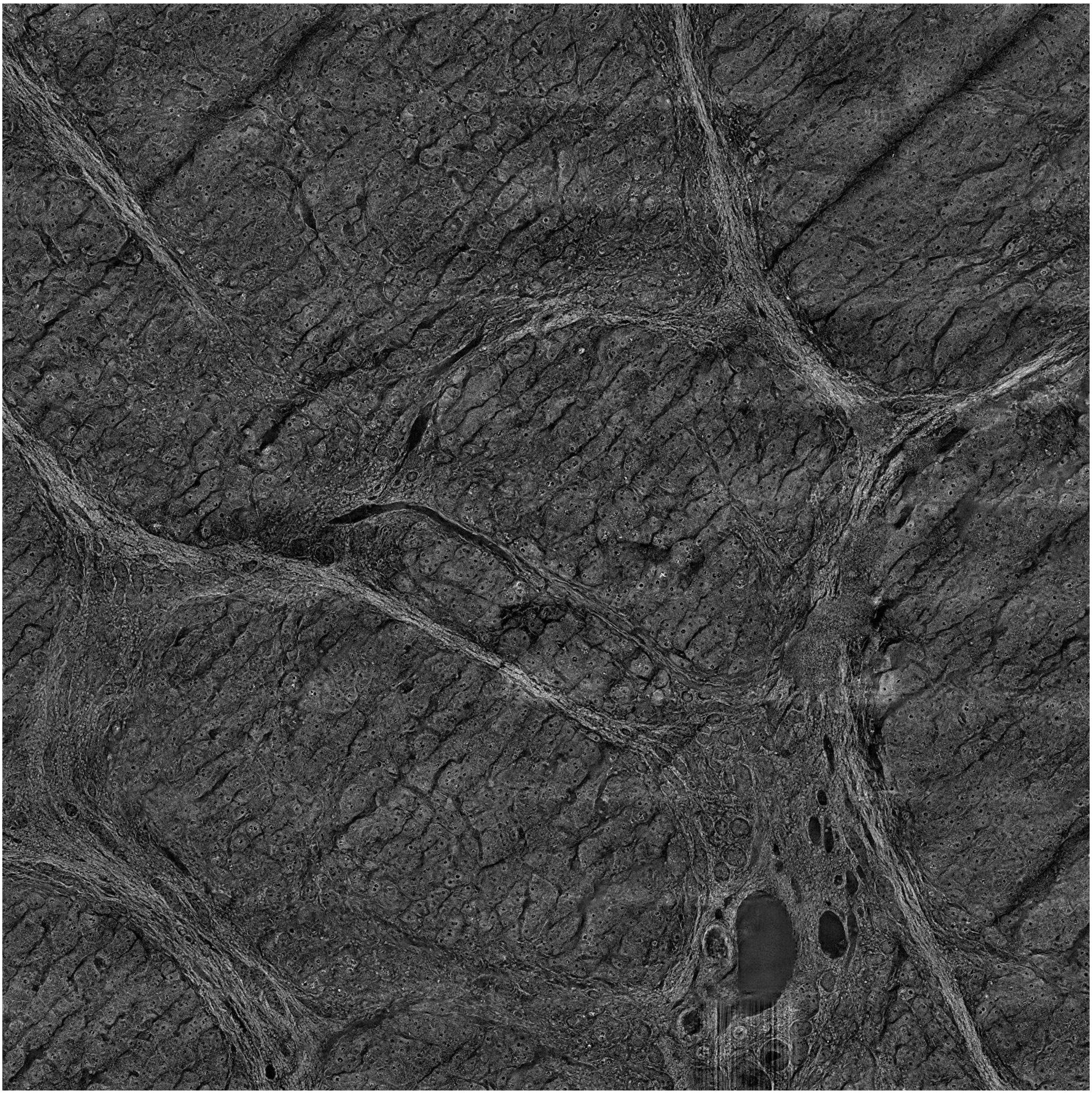

Supplementary Figure 9

Cross-sectional image of label-free 3D wide field image of the 100 μm thick tissue slices. This is the magnified version of the image shown in Fig. 3(f). The same scale and colormap used.
